# A Chairside Multimodal Platform for Temporomandibular Joint Biomechanical Assessment: Technical Evaluation and Illustrative Application

**DOI:** 10.64898/2026.07.17.738952

**Authors:** Shuchun Sun, Brooke Damon, Jichao Zhao, Konstantinia Almpani, Rachel Chung, Priyam Jani, Jonathan Mei, Ishaan Mehrotra, Cherice Hill, Farhad Ahmadi, Jian Chen, Peng Chen, Elizabeth Slate, Janice Lee, Hai Yao

**Affiliations:** Clemson-MUSC Joint Bioengineering Program, Department of Bioengineering, Clemson University, Clemson, SC 29634, USA; Department of Oral Health Sciences, Medical University of South Carolina, Charleston, SC 29425, USA; National Institute of Dental and Craniofacial Research (NIDCR), National Institutes of Health (NIH), Craniofacial Anomalies and Regeneration Section, Bethesda, MD 20892, USA; Department of Statistics, Florida State University, Tallahassee, FL, USA

**Author notes:** These authors contributed equally to this work (Shuchun Sun, Brooke Damon, Jichao Zhao). Corresponding author. Janice Lee, Address: Room 5-2531, 10 Center Dr, Bethesda, MD 20892; Hai Yao, Address: 212 Bioengineering Building, 68 President Street, Charleston, SC 29425. The authors have declared that no conflict of interest exists.

**Keywords:** Temporomandibular joint, Biomechanics, Motion capture, Electromyography, Bite force, Computational modeling

## Abstract

**Background:** Temporomandibular joint (TMJ) biomechanics can be characterized by mandibular motion, masticatory muscle activity, and bite force generation. When acquired synchronously, these functional variables can serve as model-ready inputs for subject-specific computational analyses of internal joint mechanics. However, existing tools typically measure these signals using separate hardware and software platforms, limiting synchronized acquisition within a clinically practical chairside workflow.

**Methods:** We developed and technically evaluated a compact multimodal platform for chairside acquisition of TMJ functional data and demonstrated its analytical utility in an illustrative orthognathic surgery application. The platform integrates motion, bite force, muscle activity, acoustic, and event-timing measurements with software for real-time preview, protocol guidance, and synchronized export. We assessed technical performance and chairside feasibility and analyzed representative pre- and postoperative data from an orthognathic surgery patient using kinematic, force-control, and computational modeling workflows.

**Findings:** Motion capture demonstrated submillimeter accuracy, with static and dynamic errors of approximately 0.04 mm and 0.12 mm. Bite force sensors showed excellent linearity (R² = 0.998). Chairside deployment required approximately 15 minutes each for setup and data collection. The illustrative case demonstrated that synchronized chairside data can support preoperative and postoperative kinematic analysis, bite force control capacity assessment, and estimation of TMJ disc stress.

**Interpretation:** The proposed platform enables time-efficient chairside acquisition of synchronized, model-ready multimodal datasets for quantitative TMJ biomechanical assessment. This platform and workflow could support subject-specific biomechanical analysis and future clinical studies of temporomandibular joint function.

## 1. Introduction

The temporomandibular joint (TMJ) is a paired, mechanically coupled, load-bearing joint system that coordinates mandibular rotation and translation during oral functions such as biting, chewing, and speaking (Tanaka et al., 2008; Woodford et al., 2020). During these functions, TMJ biomechanics can be characterized by complex mandibular motion, coordinated masticatory muscle activation, and bite force generation through occlusal contact. These measurable functional variables provide important information about jaw function and joint biomechanics. In addition, when acquired in a synchronized manner, they can serve as subject-specific inputs for computational models that estimate internal joint mechanical quantities that cannot be measured directly in vivo, such as joint reaction forces, articular contact conditions, and disc stresses (Sagl et al., 2019). However, generating such synchronized, model-ready functional datasets within a clinically practical chairside workflow remains challenging.

Existing technologies address portions of this workflow, but none currently provide an integrated chairside solution for synchronized acquisition of motion, loading, and muscle activity. Systems such as Planmeca (van der Helm et al., 2023) and Sicat JMT+ (Aslanidou et al., 2020) capture mandibular movement using CBCT-integrated jaw-tracking workflows, but they are not designed for chairside use and are limited in their ability to evaluate multiple oral tasks across several sessions due to the use of ionizing radiation. Other systems use techniques like magnets, infrared sensors, or optical cameras to track the trajectories of markers attached to the mandible or skull (Nagy et al., 2023; Peng et al., 2023; Woodford et al., 2020). Research laboratories have also developed customized jaw motion capture systems and emerging accelerometry-based systems with potential for future chairside or take-home use (Jucevičius et al., 2021). Other than motion capture, masticatory muscle function is commonly assessed by palpation or commercial EMG products (Wei et al., 2017), and occlusal function can be evaluated using bite force measurement devices such as those provided by Tekscan (Huang et al., 2022). Although existing technologies can measure individual components of TMJ function, and some research workflows have integrated multiple functional measurements (Gallo et al., 2015; Iwasaki et al., 2017), these approaches have generally remained lab-based. Multimodal acquisition often requires separate devices and software interfaces for jaw motion, muscle activity, and occlusal loading, leading to lengthy setup and acquisition procedures, specialized operator requirements, and post hoc data synchronization and formatting before downstream analysis. Thus, whether synchronized, model-ready TMJ functional datasets can be generated within the time, space, and personnel constraints of routine chairside workflows remains unclear.

To address these challenges, we developed a chairside multimodal platform for synchronized acquisition of TMJ functional data within a unified hardware and software workflow designed to fit the time and space constraints of routine dental examination rooms. The system integrates optical mandibular motion capture, static and dynamic bite force measurement, surface electromyography, TMJ acoustic recording, and patient-reported task-related event timing, together with software for real-time data preview, protocol guidance, and synchronized data export. By integrating hardware, real-time preview, protocol guidance, and synchronized export into a single workflow, the platform was designed to reduce multi-device operation and generate model-ready datasets at dental chairside for quantitative TMJ biomechanical assessment and downstream subject-specific modeling. In this study, we describe system design and evaluate its technical performance. We demonstrate its chairside workflow and biomechanical analytical utility in an illustrative orthognathic surgery patient through kinematic analysis, bite force control capacity assessment, and subject-specific computational modeling. This application was selected because orthognathic surgery can alter craniofacial geometry, occlusion, and the TMJ biomechanical environment (Verhelst et al., 2019). This workflow may support future larger-scale studies of patient-specific TMJ biomechanics.

## 2. Materials and Methods

The TMJ biomechanical function assessment platform consisted of compact hardware modules and integrated software for synchronized chairside data acquisition (**Figure 1**). Hardware modules captured mandibular motion, static and dynamic bite force with patient-reported event timing, surface EMG, and TMJ acoustic signals. The software provided real-time preview, protocol guidance, synchronized recording, and data export. The platform was technically benchmarked, deployed in dental examination rooms to evaluate chairside workflow feasibility, and used to generate data for kinematic analysis, bite force control capacity assessment, and subject-specific computational modeling.

**Figure 1:**
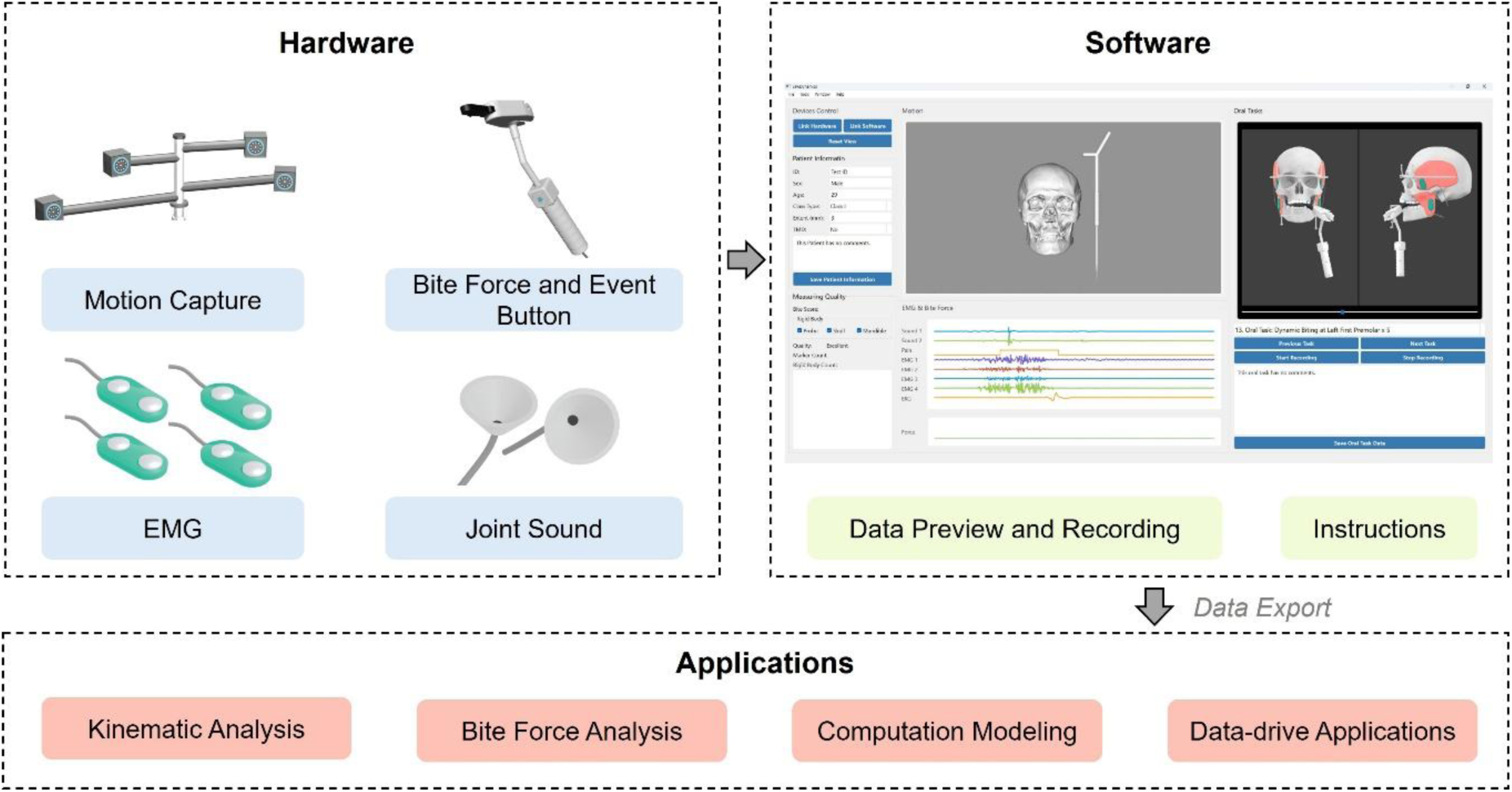
Overview of the chairside multimodal TMJ biomechanical assessment platform. The platform integrates hardware modules for mandibular motion capture, bite force measurement, surface electromyography, TMJ acoustic recording, and patient-reported event timing with software for protocol guidance, real-time preview, synchronized recording, and data export. The synchronized dataset supports downstream kinematic analysis, bite force control capacity assessment, and subject-specific computational modeling.

### 2.1. Hardware Design

The motion capture module was equipped with four high-speed infrared cameras (Prime 13, Optitrack, OR) mounted on a tripod. Cameras were calibrated before each repositioning or chairside setup using a manufacturer-provided calibration wand (OptiTrack, OR). After calibration, the camera data were used to reconstruct the 3D trajectories of 5mm reflective markers mounted on rigid frames. To ensure accurate rigid-body motion capture, seven reflective markers were assembled on a spectacle-shaped skull frame. The skull frame was worn during both motion capture and CBCT imaging, thereby enabling registration between the motion capture system and CBCT coordinates. Six reflective markers were assembled on a mandible frame. This frame was connected to the dentition via a disposable bracket attached to the buccal surface of the anterior mandibular arch using a two-part bis-acrylic polymer (Tempsmart, GC America), providing rigid coupling to the mandible and minimizing soft tissue motion artifacts. Additional setup details are provided in **Supplementary Note 1**.

The bite force and event recording module consisted of a custom-designed device incorporating static and dynamic bite force measurement components and a patient-operated event button. In static mode, the patient bit on a force sensor, enabling measurement of bite force at a nearly closed mouth position. In dynamic mode, the patient bit on two hinged bite surfaces, one containing a force sensor, with resistance provided by a torsional spring; this configuration enabled measurement of bite force during mandibular motion under loading. An event button integrated into the handle recorded patient-reported event timing for retrospective alignment with mandibular position, muscle activity, and bite force.

The surface EMG module consisted of a multichannel wired EMG system. SX230 sensors (Biometrics, Newport, UK) were placed bilaterally on the masseter and temporalis muscles to record masticatory muscle activity. The TMJ acoustic recording module consisted of a high-sensitivity MEMS microphone (CMM-3729, CUI Devices) positioned directly over the TMJ region to record acoustic signals. Analog signals from the bite force device, event button, EMG sensors, and TMJ microphone were transmitted to a custom data collection hub, which amplified and digitized all channels at 5000 Hz before streaming them to the data collection computer. All channels were synchronized through the same data acquisition system to ensure accurate temporal alignment across modalities.

### 2.2. Software Design

The software provided real-time preview, synchronized recording, protocol guidance, and data export through a unified interface (**Supplementary Figure 1**). Step-by-step examiner reminders and patient-facing instructions supported standardized task execution.

### 2.3. Benchmark Testing

To evaluate the benchmark performance of our system, we assessed both the motion capture and bite force measurement components (Furtado et al., 2013). For motion capture, a custom-made 70-mm test bar with reflective markers at both ends was measured with a micrometer (**Supplementary Figure 2**). Static accuracy was assessed by placing the bar on a table and comparing the inter-marker distance measured by our system with micrometer reference. Dynamic accuracy was evaluated by manually moving the bar for 30 seconds and calculating the root mean square error (RMSE) of the captured distance. Both static and dynamic tests were repeated five times, with the entire system reset and recalibrated between measurements, and the mean values and standard deviations were used to quantify accuracy and repeatability. To assess bite force sensor performance, known weights spanning the tested load range were applied to the sensors, and the recorded outputs were compared with the expected values. Linearity was determined by plotting applied weights against sensor readings and calculating the coefficient of determination (R^2^) from linear regression. To evaluate the synchronization between the event button and physiological signals, participants were instructed to press and hold the event button for ∼3 s during occlusion and release it when opening the mouth. Simultaneously, bilateral masseter and temporalis EMG activity and bite force were recorded, with EMG processing details provided in **Supplementary Note 2**. Synchronization was assessed by calculating the time delay between the event signal and the corresponding bite force signal at both press and release. To evaluate TMJ acoustic recording, microphone signals were acquired concurrently with mandibular motion during mouth opening and closing. Acoustic events were operationally defined as discrete, transient peaks distinguishable from baseline noise in the synchronized recording, and their corresponding timings were assigned from the acoustic trace. To assess observer agreement for sound-event identification, three observers independently reviewed the same recordings using the same standardized event criterion, and inter-observer agreement was quantified using the intraclass correlation coefficient (ICC).

### 2.4. Dental Chairside Testing and Data Analysis

For chairside testing, the TMJ biomechanical function assessment platform was installed in standard dental examination rooms at the NIH clinical center and Medical University of South Carolina. Clinical examiners underwent a 60-minute training session before collecting data. The protocol included both kinematic tasks and loaded tasks, as detailed in **Supplementary Table 1**. The total duration of each patient’s functional assessment was recorded to evaluate chairside workflow feasibility. Collected data were exported for subsequent offline analyses, including kinematic analysis, bite force control capacity evaluation, and multibody dynamic and finite element modeling. These analyses were performed in a skeletal Class III patient without clinical TMJ symptoms before and after Le Fort I osteotomy. The choice of surgical procedure was based on multiple clinical considerations and is described in **Supplementary Note 3**. The study was approved by the Institutional Review Board (IRB) of the NIH Clinical Center.

For kinematic analysis, mandible and maxilla geometries were manually segmented from CBCT images. Using the initial closed-mouth marker positions, a one-time registration was performed to align the marker frames to CBCT-derived geometries (Sorkine-Hornung and Rabinovich, 2017). Points of interest on the mandible, including the bilateral condylar apexes and the central incisor, were manually identified on the CBCT-derived geometry. The trajectories of these points were computed during mandibular motion, and relative displacements and mandibular rotation around the axis connecting the bilateral temporomandibular joints were calculated (Chen et al., 2021; Hill et al., 2023).

Bite force control capacity was assessed during the static bite task by evaluating the patient’s ability to follow a reference bite force curve. The software displayed reference levels of 10, 20, 30, 40, and 50 N, while the actual bite force was simultaneously recorded. Control capacity was quantified using the average deviation between the recorded and reference force curves, with smaller deviations indicating better force control.

The collected biomechanical functional data were combined with CBCT-derived bone morphology to construct subject-specific multibody dynamics and finite element models in ArtiSynth (Lloyd et al., 2012), used to quantify TMJ biomechanical indicators such as joint force and disc stress (Sagl et al., 2019). Forward dynamics with force tracking was employed, in which EMG and bite force measurements from the functional assessment system served as inputs to estimate TMJ joint force and disc stress (Ahmadi et al., 2026).

## 3. Results

### 3.1. Benchmark Test

The motion capture module demonstrated excellent accuracy and repeatability when evaluated against a 70-mm test bar. In static tests, the measured bar length was 69.96±0.02 mm across five trials (mean±SD, **Supplementary Table 2**). In dynamic tests, the mean measured length was 69.88±0.11 mm (mean±SD, **Supplementary Table 3**). To further assess tracking fidelity during motion, the point-to-point deviation was quantified, yielding a mean RMSE of 0.16±0.09 mm (mean±SD). The bite force sensors also showed excellent linearity: measured outputs closely matched applied loads, with linear regression yielding an R^2^ of 0.998 (**Supplementary Table 4**). Event signal synchronization testing was performed as a baseline assessment of system timing accuracy. The testing demonstrated that the patient-operated button signals were reliably aligned with bite force and EMG signals (**Figure 2**). When participants pressed and released the button, the recorded event signal consistently lagged the bite force signal by mean delay 319±177ms (mean±SD) across trials. TMJ acoustic signals were recorded during mouth opening and closing. Repeated acoustic events were identified from the microphone traces, and their corresponding timings were determined (**Supplementary Figure 3**). Using the same standardized event criterion, three observers identified the same events in the evaluated recordings, yielding excellent inter-observer agreement (ICC = 0.99).

**Figure 2:**
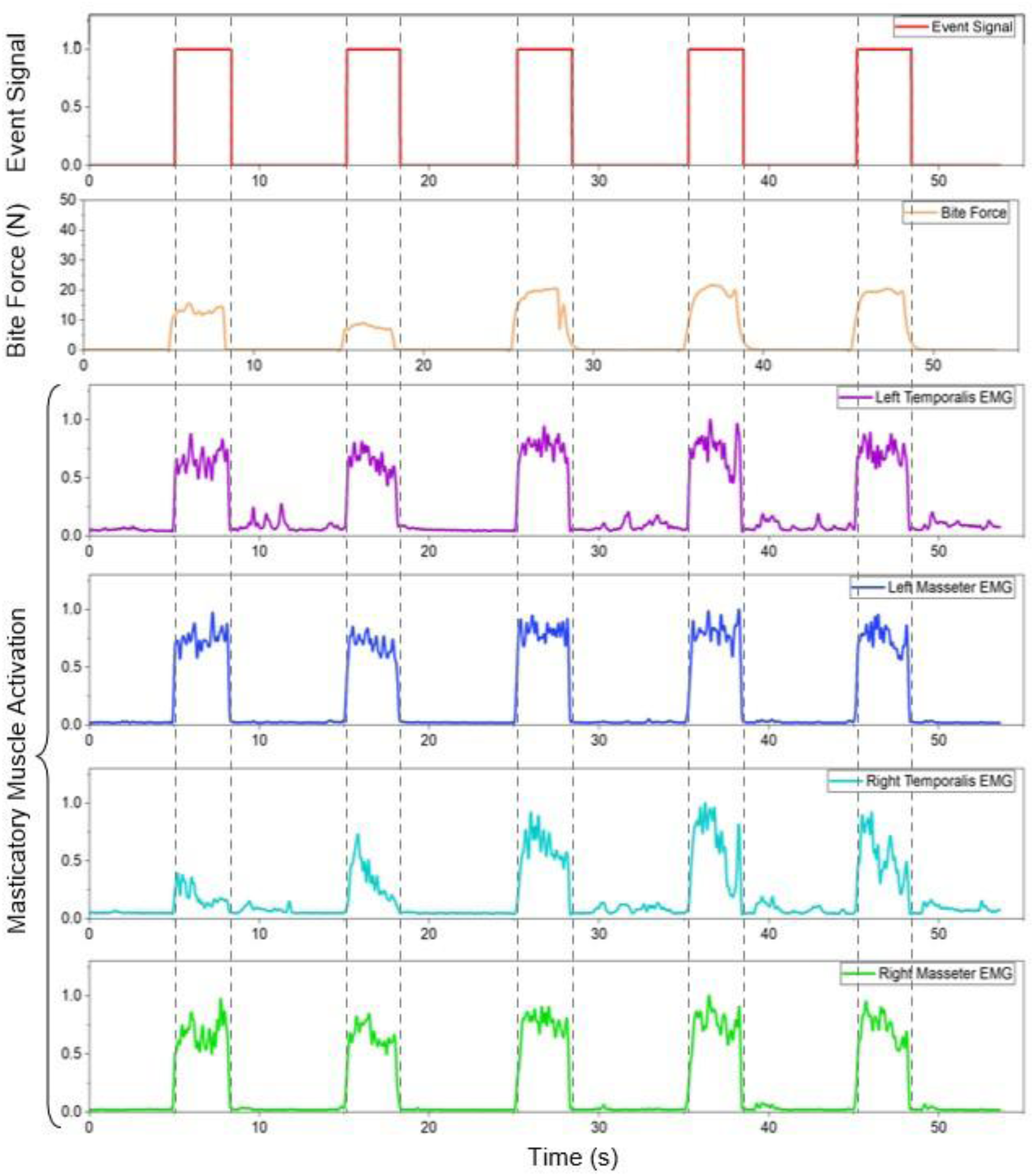
Event signal synchronization testing demonstrated that the patient-operated button signals were reliably aligned with bite force and EMG signals.

### 3.2. Dental Chairside Test

Our biomechanical functional system was successfully deployed in dental examination rooms. The compact design allows all components to fit into two small suitcases (∼22×14×9 inches) with a total weight under 10 kg, ensuring easy portability. The system fit comfortably within a standard dental examination room (3.2×3.0 m) without interfering with routine equipment. Clinical examiners, after a 60-minute training, were able to operate the system efficiently. Setup and calibration required ∼15 minutes, and each data collection session was completed in another ∼15 minutes. For the illustrative skeletal Class III case, no patient-reported task-related pain or acoustic event was observed in either the preoperative or postoperative assessment. The representative case results therefore focus on kinematic, bite force control capacity, and biomechanical modeling outcomes.

Kinematic analysis was performed for maximum open-close motion before and after the Le Fort I surgery (**Figure 3A**). In this analysis, incisor displacement was comparable pre- and postoperatively, with maximum opening of 39.00 mm before surgery and 36.29 mm after surgery (**Figure 3B**). In contrast, TMJ apex displacement decreased substantially after surgery on both sides. The maximum TMJ apex displacement decreased from 10.38 mm to 4.85 mm on the left (**Figure 3C**) and from 9.76 mm to 5.43 mm on the right (**Figure 3D**). Despite similar mouth opening, the joint apex kinematic indicator suggested increased rotation and decreased translation postoperatively.

**Figure 3:**
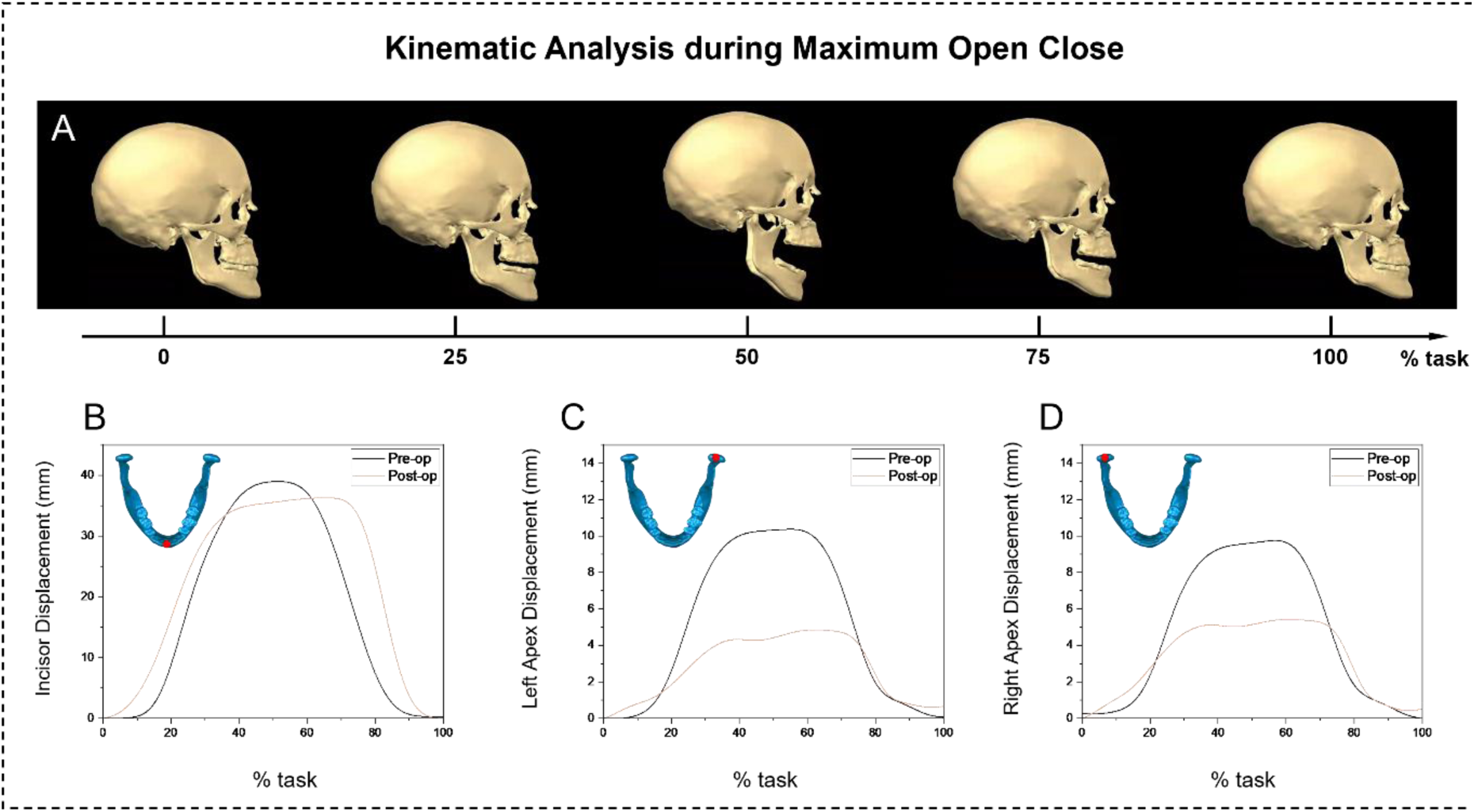
Kinematic analysis before and after Le Fort I surgery.

In this illustrative case, target-force tracking deviation was lower postoperatively. Preoperatively, the average deviation from the reference bite force profile was 8.97 N at the left premolar and 7.07 N at the right premolar, indicating limited control. Postoperatively, the deviations decreased to 3.91 N and 4.22 N, respectively. Because smaller deviations reflect closer tracking of the reference curve, these results demonstrate how the platform can quantify bite force control capacity across pre- and postoperative assessments (**Figure 4**).

**Figure 4:**
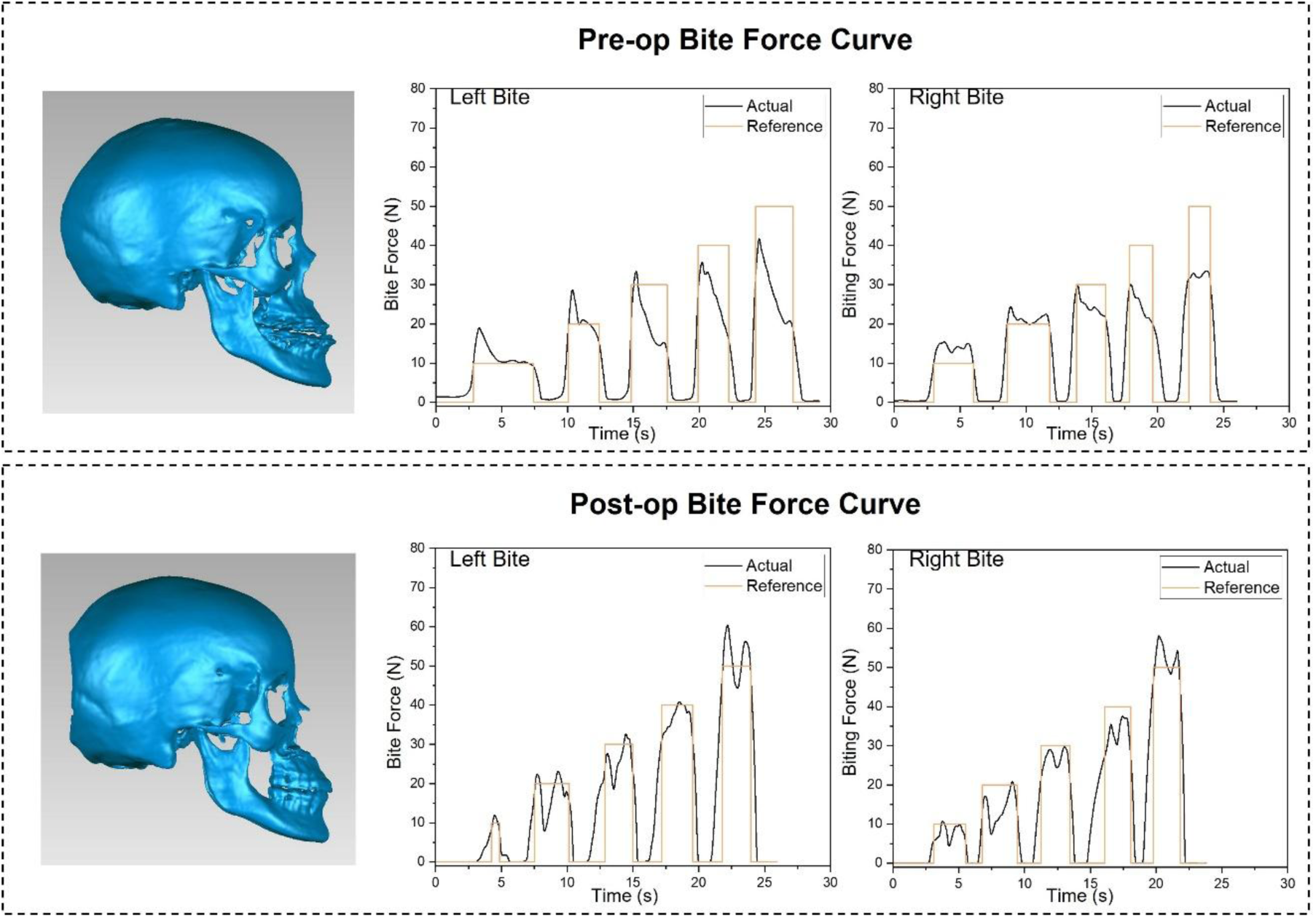
Bite force control capacity assessed by target-force tracking at the left and right premolars before and after Le Fort I surgery.

Computational models were built for the Class III subject in both preoperative and postoperative states. The simulations were driven by measured EMG activity together with approximately 20 N bite force recorded by our system, and outputs included TMJ disc stresses. During left premolar biting, stress in the left TMJ disc decreased, whereas stress in the right disc increased. During right premolar biting, stress in both the left and right TMJ discs decreased. Detailed maximum stress values are provided in **Supplementary Table 5**. However, the magnitude of these model-estimated stress differences was small in this illustrative analysis (**Figure 5**).

**Figure 5:**
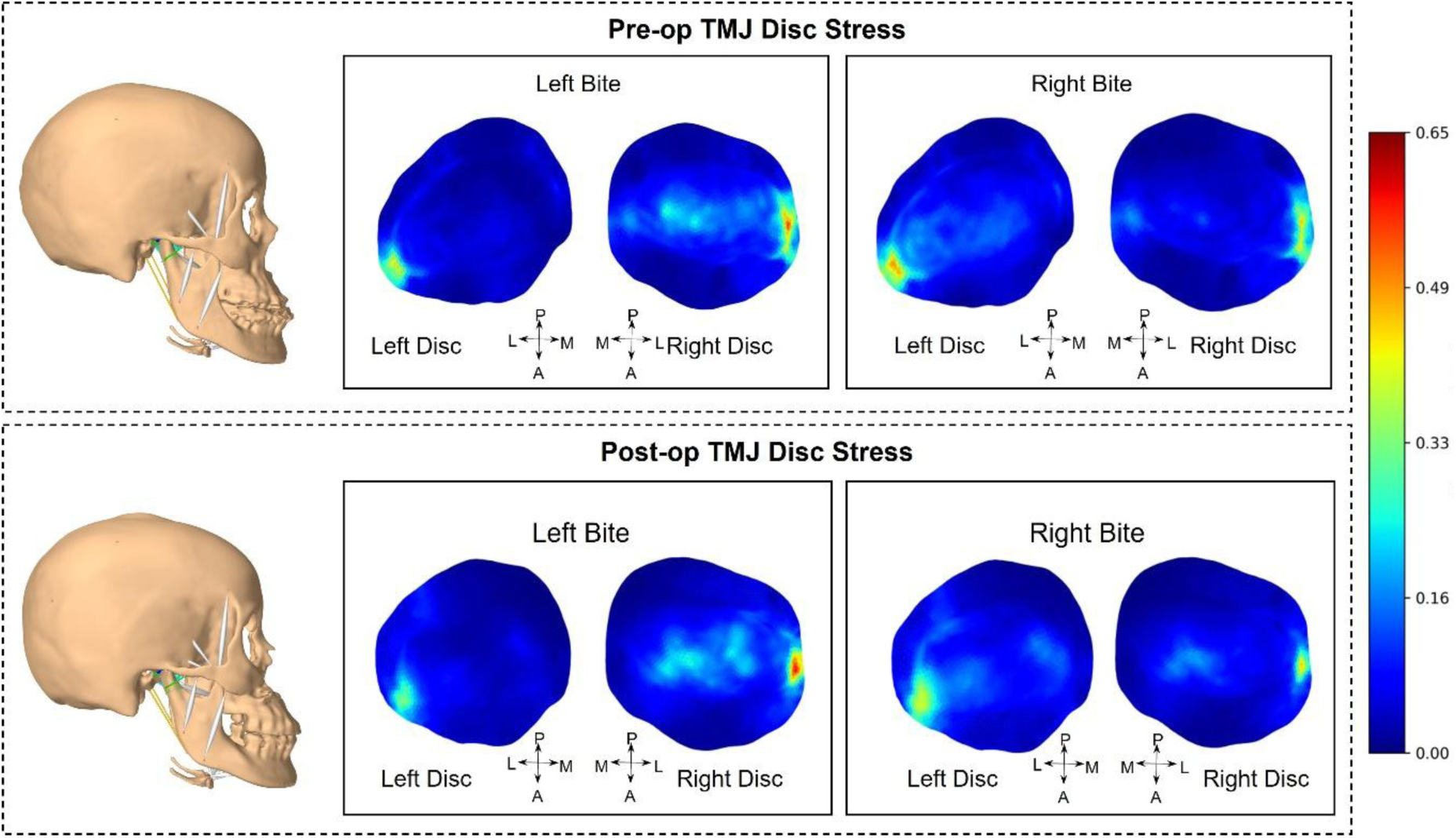
Multibody dynamic and finite element modeling results of TMJ disc stress during premolar biting before and after Le Fort I surgery.

## 4. Discussion

In this study, we developed and benchmarked a compact, multimodal chairside system for integrated assessment of TMJ biomechanics. Benchmark testing confirmed high accuracy and repeatability of the motion capture and bite force modules, and chairside deployment demonstrated the system’s feasibility, portability, and ease of use in routine dental examination rooms. Through an illustrative case study, we further showed how the system enables subject-specific analyses of mandibular kinematics, bite force control capacity, and computational modeling of TMJ joint loading and disc stress. Collectively, these findings highlight the potential of the system to generate synchronized, model-ready multimodal datasets for quantitative and subject-specific TMJ biomechanical assessment in chairside and research settings.

The benchmark test results confirmed the performance of our TMJ biomechanical function assessment platform. Since TMJ movements are inherently small (Lendraitiene et al., 2021), high precision of joint motion capture is required for clinical and research utility. Our results demonstrated that our system achieved comparable or superior accuracy relative to previously reported jaw tracking systems. For example, prior studies of optical and magnetic jaw tracking systems have reported static errors ranging from 0.03 to 0.80 mm, and dynamic errors up to ∼1.0 mm depending on the system and task complexity (Baltali et al., 2008; Coutant et al., 2008; Sadat-Khonsari et al., 2003; Woodford et al., 2020). In contrast, our system maintained a static accuracy of 0.04 mm and a dynamic accuracy of 0.12 mm, underscoring its suitability for precise TMJ motion analysis. The synchronized EMG and bite force recordings further support integrated functional measurement and downstream computational modeling. Inaccurate functional inputs can compromise estimates of muscle forces and joint loads (Beek et al., 2000; Lloyd et al., 2012). Event signal synchronization testing further supported the system’s ability to time-stamp patient-reported task-related events and align them with objective biomechanical measurements. The observed ∼300ms delay between the event button and bite force signals is consistent with expected human sensorimotor response times (Woods et al., 2015). This indicates that the recorded delay largely reflects participant response latency rather than system-level lag. The acoustic channel captured transient sound events during mandibular motion, and their timings could be assigned from the synchronized acoustic trace for retrospective alignment with mandibular kinematics, EMG, bite force, and patient-reported event signals. In this study, acoustic events were identified using a standardized visual criterion based on discrete, transient peaks distinguishable from baseline noise, and observer agreement under this criterion was excellent. However, no predefined amplitude threshold, frequency-domain criterion, or automated detection algorithm was applied. Therefore, these annotations should be interpreted as a technical demonstration of synchronized acoustic recording rather than validation of pathological TMJ click detection.

Dental chairside testing further demonstrated the system’s practicality in a real clinical environment. In contrast to laboratory-based multimodal workflows that often require specialized personnel, extended setup and acquisition time, and coordination across multiple devices or software interfaces (Gallo et al., 2015; Iwasaki et al., 2017), the present platform was operated by trained clinical examiners and completed within a chairside workflow. Setup and calibration required approximately 15 minutes, and functional assessments across multiple oral tasks were completed in another approximately 15 minutes. The collected data illustrated multiple biomechanical outputs. For example, kinematic analyses captured mandibular and condylar motion patterns, bite force control capacity assessment quantified deviation from target force profiles, and multibody dynamic and finite element models provided estimates of TMJ disc stresses. Such synchronized datasets open the possibility for applying advanced techniques that synthesize information across multimodal functional data, such as computational modeling (She et al., 2021) and machine learning (Sun et al., 2024), to identify biomechanical patterns across patients, tasks, and treatment states. In addition, the patient-operated event button enables task-related patient-reported events, including pain onset in future symptomatic applications, to be time-stamped and retrospectively linked to the corresponding biomechanical state. Because the illustrative orthognathic surgery case did not exhibit patient-reported task-related pain or acoustic events, the representative analyses focused on kinematic, bite force control capacity, and computational modeling outcomes. Present analyses serve as proof-of-concept demonstrations of how the system can be used to analyze coordinated biomechanical changes in a paired pre- and postoperative setting. The small model-estimated stress differences are also plausible because Le Fort I osteotomy primarily alters the maxilla rather than directly repositioning the mandible. Because advanced kinematic and computational modeling analyses require customized processing and quality control, these analyses were performed offline rather than in real time. Future software development could streamline selected analysis modules for more automated chairside implementation.

Despite its promising capabilities, several limitations should be noted. First, the orthognathic surgery analysis was an illustrative single-case application and was not intended to establish clinical validity, treatment-related effects, or population-level biomechanical conclusions. Second, because the case did not exhibit patient-reported task-related pain or acoustic events, the event and acoustic channels were demonstrated primarily as synchronized recording capabilities rather than validated clinical outcome measures. Finally, advanced kinematic and computational modeling analyses currently require data export and customized offline processing, which limits immediate in-clinic usability. Future software development should focus on streamlining selected analysis modules for larger-scale chairside studies.

## 5. Conclusion

The proposed TMJ biomechanical function assessment platform enables accurate, user-friendly, and integrated chairside acquisition of multimodal TMJ functional data. Benchmark testing confirmed its accuracy and reliability, while chairside deployment demonstrated its practicality in dental settings. The data analysis examples demonstrated that synchronized chairside data can support downstream kinematic analysis, bite force control capacity assessment, and subject-specific computational modeling of TMJ disc stress. Future work will focus on streamlining analysis modules and evaluating the platform in larger cohorts.

## Clinical Trial Number

Clinical trial number: not applicable

## CRediT authorship contribution statement

**Shuchun Sun:** Conceptualization, Software, Formal analysis, Data curation, Visualization, Writing – original draft, Writing – review & editing. **Brooke Damon:** Conceptualization, Methodology, Investigation, Writing – review & editing. **Jichao Zhao:** Software, Formal analysis, Visualization, Writing – original draft, Writing – review & editing. **Konstantinia Almpani:** Data curation, Investigation, Writing – review & editing. **Rachel Chung:** Data curation, Investigation, Writing – review & editing. **Priyam Jani:** Data curation, Investigation, Writing – review & editing. **Jonathan Mei:** Formal analysis, Writing – review & editing. **Ishaan Mehrotra:** Investigation, Writing – review & editing. **Cherice Hill:** Formal analysis, Writing – review & editing. **Farhad Ahmadi:** Formal analysis, Writing – review & editing. **Jian Chen:** Visualization, Writing – original draft, Writing – review & editing. **Peng Chen:** Data curation, Formal analysis, Writing – original draft, Writing – review & editing. **Elizabeth Slate:** Data curation, Formal analysis, Writing – original draft, Writing – review & editing. **Janice Lee:** Conceptualization, Funding acquisition, Supervision, Writing – review & editing. **Hai Yao:** Conceptualization, Funding acquisition, Supervision, Formal analysis, Writing – original draft, Writing – review & editing.

## Statements and Declarations

The contributions of the NIH authors were made as part of their official duties as NIH federal employees, are in compliance with agency policy requirements, and are considered Works of the United States Government. However, the findings and conclusions presented in this paper are those of the authors and do not necessarily reflect the views of the NIH or the U.S. Department of Health and Human Services.

## Data availability

Benchmark testing results are included in the supplementary materials. The input data and results for the kinematic analysis, bite force analysis, and computational modeling are available from the corresponding author upon reasonable request.

## References

Ahmadi, F., Sun, S., Zhao, J., Chen, J., Wilson, M.B., Damon, B., Wu, Y., Almpani, K., Chung, R., Jani, P., Chen, P., Slate, E.H., Lee, J.S., Sagl, B., Yao, H., 2026. A Computational Framework for Simulating Patient-Specific TMJ Biomechanics Using a Combined Multibody Dynamics and Finite Element Approach. Ann Biomed Eng. doi: 10.1007/s10439-026-04020-0

Aslanidou, K., Xie, R., Christou, T., Lamani, E., Kau, C.H., 2020. Evaluation of temporomandibular joint function after orthognathic surgery using a jaw tracker. J Orthod 47, 140–148. doi: 10.1177/1465312520908277

Baltali, E., Zhao, K.D., Koff, M.F., Durmuş, E., An, K.-N., Keller, E.E., 2008. A method for quantifying condylar motion in patients with osteoarthritis using an electromagnetic tracking device and computed tomography imaging. J Oral Maxillofac Surg 66, 848–857. doi: 10.1016/j.joms.2008.01.021

Beek, M., Koolstra, J.H., van Ruijven, L.J., van Eijden, T.M., 2000. Three-dimensional finite element analysis of the human temporomandibular joint disc. J Biomech 33, 307–316. doi: 10.1016/s0021-9290(99)00168-2

Chen, C.C., Lin, C.C., Hsieh, H.P., Fu, Y.C., Chen, Y.J., Lu, T.W., 2021. In vivo three-dimensional mandibular kinematics and functional point trajectories during temporomandibular activities using 3d fluoroscopy. Dentomaxillofac Radiol 50, 20190464. doi: 10.1259/dmfr.20190464

Coutant, J.-C., Mesnard, M., Morlier, J., Ballu, A., Cid, M., 2008. Discrimination of objective kinematic characters in temporomandibular joint displacements. Arch Oral Biol 53, 453–461. doi: 10.1016/j.archoralbio.2007.11.010

Furtado, D.A., Pereira, A.A., Andrade Ade, O., Bellomo, D.P., Jr., da Silva, M.R., 2013. A specialized motion capture system for real-time analysis of mandibular movements using infrared cameras. Biomed Eng Online 12, 17. doi: 10.1186/1475-925x-12-17

Gallo, L., Iwasaki, L., Gonzalez, Y., Liu, H., Marx, D., Nickel, J., 2015. Diagnostic group differences in temporomandibular joint energy densities. Orthodontics & craniofacial research 18, 164–169

Hill, C., Sun, S., Mei, A., Almpani, K., Jani, P., Ahmadi, F., Damon, B., Slate, E., Yao, H., Lee, J.S., 2023. Normative 3D Mandibular and TMJ Kinematics in Orthognathic Asymptomatic and Symptomatic Patients. Journal of Oral and Maxillofacial Surgery 81, S5–S7. doi: 10.1016/j.joms.2023.08.075

Huang, Y.F., Wang, C.M., Shieh, W.Y., Liao, Y.F., Hong, H.H., Chang, C.T., 2022. The correlation between two occlusal analyzers for the measurement of bite force. BMC Oral Health 22, 472. doi: 10.1186/s12903-022-02484-9

Iwasaki, L., Gonzalez, Y., Liu, Y., Liu, H., Markova, M., Gallo, L., Nickel, J., 2017. Mechanobehavioral scores in women with and without TMJ disc displacement. Journal of dental research 96, 895–901

Jucevičius, M., Ožiūnas, R., Mažeika, M., Marozas, V., Jegelevičius, D., 2021. Accelerometry-Enhanced Magnetic Sensor for Intra-Oral Continuous Jaw Motion Tracking. Sensors (Basel) 21. doi: 10.3390/s21041409

Lendraitiene, E., Smilgiene, L., Petruseviciene, D., Savickas, R., 2021. Changes and Associations between Cervical Range of Motion, Pain, Temporomandibular Joint Range of Motion and Quality of Life in Individuals with Migraine Applying Physiotherapy: A Pilot Study. Medicina (Kaunas) 57, 630. doi: 10.3390/medicina57060630

Lloyd, J.E., Stavness, I., Fels, S., 2012. ArtiSynth: A fast interactive biomechanical modeling toolkit combining multibody and finite element simulation, Soft tissue biomechanical modeling for computer assisted surgery. Springer, pp. 355–394.

Nagy, Z., Mikolicz, A., Vag, J., 2023. In-vitro accuracy of a novel jaw-tracking technology. J Dent 138, 104730. doi: 10.1016/j.jdent.2023.104730

Peng, T., Yang, Z., Ma, T., Zhang, M., Ren, G., 2023. Comparative evaluation of the volume of occlusal adjustment of repositioning occlusal devices designed by using an average value digital articulator and the jaw movement analyzer. J Prosthet Dent. doi: 10.1016/j.prosdent.2023.06.018

Sadat-Khonsari, R., Fenske, C., Kahl-Nieke, B., Kirsch, I., Jüde, H.D., 2003. Mandibular instantaneous centers of rotation in patients with and without temporomandibular dysfunction. J Orofac Orthop 64, 256–264. doi: 10.1007/s00056-003-0204-z

Sagl, B., Schmid-Schwap, M., Piehslinger, E., Kundi, M., Stavness, I., 2019. A Dynamic Jaw Model With a Finite-Element Temporomandibular Joint. Front Physiol 10, 1156. doi: 10.3389/fphys.2019.01156

She, X., Sun, S., Damon, B.J., Hill, C.N., Coombs, M.C., Wei, F., Lecholop, M.K., Steed, M.B., Bacro, T.H., Slate, E.H., Zheng, N., Lee, J.S., Yao, H., 2021. Sexual dimorphisms in three-dimensional masticatory muscle attachment morphometry regulates temporomandibular joint mechanics. J Biomech 126, 110623. doi: 10.1016/j.jbiomech.2021.110623

Sorkine-Hornung, O., Rabinovich, M., 2017. Least-squares rigid motion using svd. Computing 1, 1–5

Sun, S., Xu, P., Buchweitz, N., Hill, C.N., Ahmadi, F., Wilson, M.B., Mei, A., She, X., Sagl, B., Slate, E.H., Lee, J.S., Wu, Y., Yao, H., 2024. Explainable deep learning and biomechanical modeling for TMJ disorder morphological risk factors. JCI Insight 9. doi: 10.1172/jci.insight.178578

Tanaka, E., Detamore, M., Mercuri, L., 2008. Degenerative disorders of the temporomandibular joint: etiology, diagnosis, and treatment. Journal of dental research 87, 296–307

van der Helm, H.C., Dieters, A.J., Dijkstra, P.U., van der Meer, W.J., Kuijpers-Jagtman, A.M., 2023. Exploring the Validity of an Optoelectronic Integrated Cone Beam Computed Tomography Jaw Tracking System. Journal of clinical medicine 12, 4145

Verhelst, P.J., Van der Cruyssen, F., De Laat, A., Jacobs, R., Politis, C., 2019. The Biomechanical Effect of the Sagittal Split Ramus Osteotomy on the Temporomandibular Joint: Current Perspectives on the Remodeling Spectrum. Front Physiol 10, 1021. doi: 10.3389/fphys.2019.01021

Wei, F., Van Horn, M.H., Coombs, M.C., She, X., Gonzales, T.S., Gonzalez, Y.M., Scott, J.M., Iwasaki, L.R., Nickel, J.C., Yao, H., 2017. A pilot study of nocturnal temporalis muscle activity in TMD diagnostic groups of women. J Oral Rehabil 44, 517–525. doi: 10.1111/joor.12517

Woodford, S.C., Robinson, D.L., Mehl, A., Lee, P.V.S., Ackland, D.C., 2020. Measurement of normal and pathological mandibular and temporomandibular joint kinematics: A systematic review. J Biomech 111, 109994. doi: 10.1016/j.jbiomech.2020.109994

Woods, D.L., Wyma, J.M., Yund, E.W., Herron, T.J., Reed, B., 2015. Factors influencing the latency of simple reaction time. Front Hum Neurosci 9, 131. doi: 10.3389/fnhum.2015.00131

