## Supplementary Materials for "A Chairside Multimodal Platform for Temporomandibular Joint Biomechanical Assessment: Technical Evaluation and Illustrative Application"

*Corresponding author.

Janice Lee:

Address: Room 5-2531, 10 Center Dr, Bethesda, MD 20892

Hai Yao:

Address: 212 Bioengineering Building, 68 President Street, Charleston, SC 29425

**Table of Content**

**Supplementary Figure 1:** Software modules of the system.

**Supplementary Figure 2:** 70mm test bar with reflective markers.

**Supplementary Figure 3:** Representative TMJ acoustic recording during mouth opening and closing.

**Supplementary Table 1:** Oral tasks performed during functional assessment.

**Supplementary Table 2:** Static length measurement for five reset and recalibration.

**Supplementary Table 3:** Dynamic length measurement for five reset and recalibration.

**Supplementary Table 4:** Sensor linearity validation.

**Supplementary Table 5:** Maximum TMJ disc stress from ArtiSynth model.

**Supplementary Note 1:** Motion capture setup and calibration details.

**Supplementary Note 2:** EMG processing for event synchronization testing.

**Supplementary Note 3:** Clinical considerations for choice of surgical procedure.

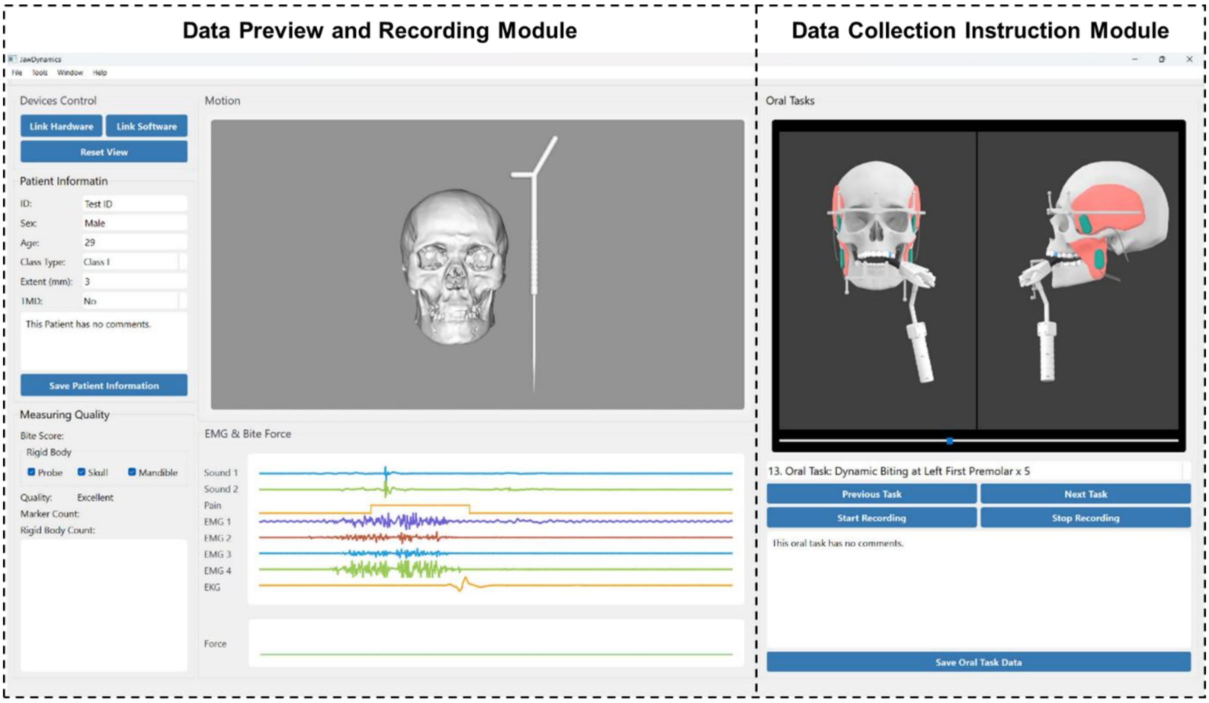


**Supplementary Figure 1:** Software modules of the system, showing (A) real-time data preview and recording and (B) system setup and data collection instructions.

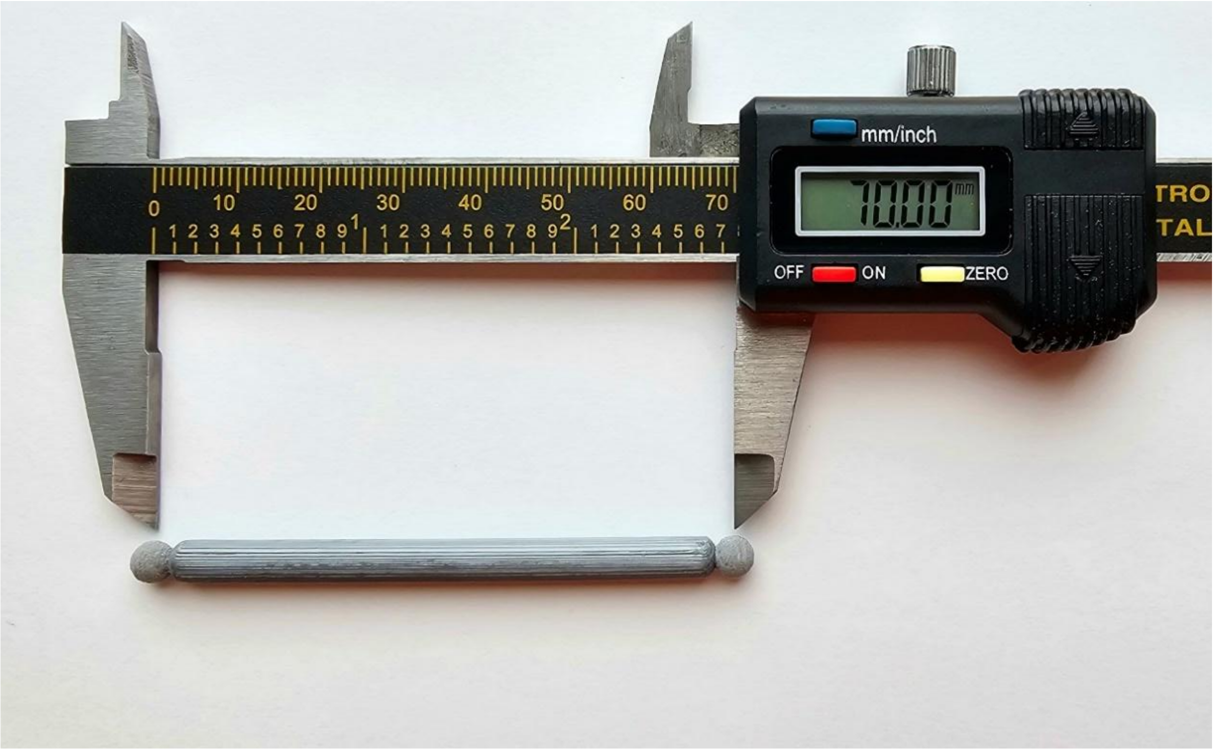


**Supplementary Figure 2:** 70mm test bar with reflective markers at both ends for motion capture benchmark testing.

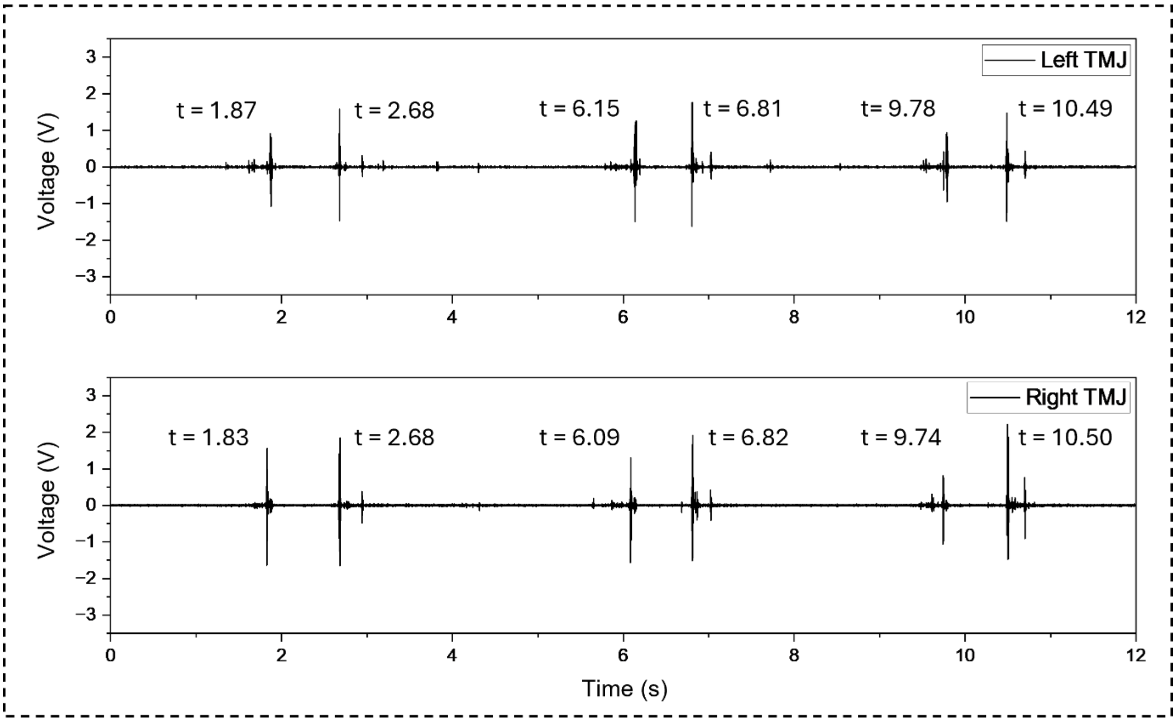


**Supplementary Figure 3:** Representative TMJ acoustic recording during mouth opening and closing. Acoustic events were identified as discrete, transient peaks distinguishable from baseline noise in the synchronized acoustic trace, and their corresponding timings were annotated.

**Supplementary Table 1**: Oral tasks performed during functional assessment.

| Step | Oral Task |
|---|---|
| 1 | Maximum Open and Close × 5 |
| 2 | Maximum Lateral Movement to the Left × 5 |
| 3 | Maximum Lateral Movement to the Right × 5 |
| 4 | Maximum Anterior Movement × 5 |
| 5 | Maximum Posterior Movement × 5 |
| 6 | Static Biting at the Left First Premolar |
| 7 | Static Biting at the Right First Premolar |
| 8 | Static Biting at the Central Incisor |
| 9 | Dynamic Biting at the Left First Premolar × 5 |
| 10 | Dynamic Biting at the Right First Premolar × 5 |
| 11 | Dynamic Biting at the Central Incisor × 5 |

**Supplementary Table 2**: Static length measurement for five reset and recalibration.

| Test 1 | Test 2 | Test 3 | Test 4 | Test 5 | Mean | Std Dev |
|---|---|---|---|---|---|---|
| 69.99mm | 69.94mm | 70.01mm | 69.98mm | 69.95mm | 69.96mm | 0.02mm |

**Supplementary Table 3**: Dynamic length measurement for five reset and recalibration.

| | **Average for 30 Secs** | **RMSE** |
|---|---|---|
| **Test 1** | 70.02 | 0.07 |
| **Test 2** | 69.86 | 0.14 |
| **Test 3** | 69.68 | 0.32 |
| **Test 4** | 69.90 | 0.18 |
| **Test 5** | 69.95 | 0.08 |
| **Mean** | 69.88 | 0.16 |
| **Std Dev** | 0.11 | 0.09 |

**Supplementary Table 4**: Sensor linearity validation.

| Force (N) | Voltage (V) |
|---|---|
| 0 | 0.0262 |
| 10 | 0.5288 |
| 20 | 0.9438 |
| 30 | 1.3813 |
| 40 | 1.7689 |
| 50 | 2.1573 |

**Supplementary Table 5**: Maximum TMJ disc stress from ArtiSynth model.

| | **Left Bite Left Disc (MPa)** | **Left Bite Right Disc (MPa)** | **Right Bite Left Disc (MPa)** | **Right Bite Right Disc (MPa)** |
|---|---|---|---|---|
| Before Surgery | 0.43 | 0.58 | 0.52 | 0.53 |
| After Surgery | 0.40 | 0.65 | 0.49 | 0.48 |

**Supplementary Note 1:** Motion capture setup and calibration details

The motion capture cameras were positioned approximately 1 m from the subject, providing sufficient space for examiner movement while maintaining tracking accuracy. The cameras were arranged in a non-collinear configuration to reduce capture errors. Prior to data collection, the camera system was calibrated using a manufacturer-provided calibration wand with a known geometric configuration. During calibration, the wand was moved throughout the capture volume, and the system reconstructed the relative positions and orientations of the cameras to establish a unified global coordinate system. The calibration procedure was performed within OptiTrack Motive software. Calibration remained valid as long as the relative camera positions were unchanged; therefore, because the system was designed for portable chairside deployment, recalibration was performed whenever the system was repositioned or set up in a new clinical environment.

The spectacle-shaped skull frame was tightly fitted using an adjustable eyeglass structure and positioned in a standardized manner for each subject. The adjustable design accommodated inter-subject variability in head size and facial anatomy while maintaining consistent frame positioning across sessions. Because the frame was supported by the nasal bridge and ears, it remained stable during mandibular motion and was minimally affected by soft tissue deformation during jaw opening and biting tasks. To accommodate variability in mandibular size and dental arch curvature, a set of mandibular attachment brackets with different geometries was designed and selected by the operator during setup.

**Supplementary Note 2:** EMG processing for event synchronization testing

The raw EMG signals recorded during event synchronization testing were high-pass filtered using a zero-lag fourth-order Butterworth filter at 30 Hz to remove DC offsets and electrode motion artifacts. The signals were then full-wave rectified and normalized to the peak rectified value. The processed signals were subsequently low-pass filtered using a zero-lag sixth-order Butterworth filter at 6 Hz to obtain the EMG envelope while attenuating high-frequency noise. Because the occlusion task involved minimal jaw displacement, kinematic data were not included in this synchronization assessment.

**Supplementary Note 3**: Clinical considerations for choice of surgical procedure

The choice of surgical procedure was based on comprehensive clinical planning. Considerations included:

(1) patient-specific esthetic considerations, including the patient's desired facial profile, preference for preservation of midfacial fullness, and broader soft tissue/facial harmony considerations in the context of the patient's background and stated esthetic goals.

(2) airway-related considerations, including the potential risk that mandibular setback could further restrict the airway, the absence of an obviously constricted airway at present, uncertainty regarding snoring history, and the possible increased long-term risk of airway compromise with aging in the setting of elevated BMI (BMI = 31.3).

(3) facial soft tissue and profile considerations, including the presence of infraorbital and midfacial flatness reflecting a combination of skeletal and soft tissue characteristics, as well as a relatively thick soft tissue biotype that was expected to mask underlying skeletal alterations to a greater extent.

(4) the goal of selecting the most conservative surgical procedure capable of achieving satisfactory functional and esthetic outcomes, particularly because the magnitude of the discrepancy was relatively small and could be addressed with single-jaw maxillary surgery.

In this case, after consideration of these factors, isolated Le Fort I osteotomy was selected as a conservative surgical approach capable of achieving satisfactory functional and esthetic outcomes for this patient.
